# Modelling the *Wolbachia* Incompatible Insect Technique: strategies for effective mosquito population elimination

**DOI:** 10.1101/2020.07.13.201483

**Authors:** D.E. Pagendam, B.J. Trewin, N. Snoad, S.A. Ritchie, A.A. Hoffmann, K.M. Staunton, C. Paton, N. Beebe

**Affiliations:** CSIRO Data61, 41 Boggo Road, Dutton Park, QLD 4102 Australia; CSIRO Health and Biosecurity, 41 Boggo Road, Dutton Park, QLD 4102 Australia; Verily Life Sciences, 259 East Grand Avenue, South San Francisco, CA 94080, USA; College of Public Health, Medical and Veterinary Sciences, James Cook University, Smithfield QLD 4878, Australia; Australian Institute of Tropical Health and Medicine, James Cook University, Smithfield QLD 4878, Australia; School of Biological Sciences, Bio21 Institute, The University of Melbourne, Parkville Vic, Australia; School of Biological Sciences, University of Queensland, St Lucia, QLD 4072, Australia

**Keywords:** incompatible insect technique, *Wolbachia*, establishment risk, elimination, stochastic model, simulation

## Abstract

The *Wolbachia* Incompatible Insect Technique (IIT) shows promise as a method for eliminating invasive mosquitoes such as *Aedes aegypti* (Linnaeus)(Diptera: Culicidae) and reducing the incidence of vector-borne diseases such as dengue, chikungunya and Zika. Successful implementation of this biological control strategy relies on high-fidelity separation of male from female insects in mass production systems for inundative release into landscapes. Processes for sex-separating mosquitoes are typically error prone, laborious and IIT programs run the risk of releasing *Wolbachia* infected females and replacing wild mosquito populations. We introduce a simple Markov Population Process (MPP) model for studying mosquito populations subjected to a *Wolbachia*-IIT program which exhibit an unstable equilibrium threshold. The model is used to study, *in silico*, scenarios that are likely to yield a successful elimination result. Our results suggest that elimination is best achieved by releasing males at rates that adapt to the ever-decreasing wild population, thus reducing the risk of releasing *Wolbachia*-infected females while reducing costs. While very high-fidelity sex-separation is required to avoid establishment, release programs tend to be robust to the release of a small number of *Wolbachia*-infected females. These findings will inform and enhance the next generation of *Wolbachia*-IIT control strategies that are already showing great promise in field trials.

## Introduction

Since the 1950s, the sterile insect technique (SIT) has been a popular biological control method for the management of pest insect populations (Dyck et al., 2006). The SIT takes advantage of the reproductive biology of a species to suppress or eliminate populations, applied through the inundative release of sterile individuals. Traditional approaches using radiation and chemosterilants have succeeded in the elimination of a variety of insect pests, including the new world screwworm in the Americas and Libya (Vargas-Terán et al., 2005), the tsetse fly in Zanzibar (Vreysen et al., 2000) and the Mediterranean fruit fly in Mexico and Guatemala (Hendrichs et al., 1983). In a modern context, most SIT programs now form a component of integrated pest management programmes, generating billions of dollars in savings for agricultural commerce annually, through the reduced environmental impacts of pesticide use (Dyck et al., 2006). The knowledge gained from these historical campaigns now provides the baseline information for a renewed interest in area-wide SIT control programs.

Rapid advances in fields such as molecular biology, genetics and computer science have seen a resurgence in SIT, particularly in attempting to prevent mosquito borne diseases such as dengue, chikungunya, Zika and yellow fever (Gilles et al., 2014, Kittayapong et al., 2018). The incompatible insect technique (IIT) is closely related to SIT, but rather than releasing sterilized insects, this approach relies upon *Wolbachia*-infected male mosquitoes that are incapable of producing viable offspring after mating with a wild-type female (Laven, 1967). Endosymbiotic bacteria, such as *Wolbachia*, may exhibit a biological mechanism known as cytoplasmic incompatibility (CI; Laven, 1967). When CI occurs, male mosquitoes infected with *Wolbachia* are not sterile *per se*, since they still produce offspring when mated with *Wolbachia*-infected females. However, in crosses with wild-type females not containing *Wolbachia* (or a compatible strain of *Wolbachia)* mating fails to produce viable offspring as eggs fail to hatch.

Sterile mosquito strategies typically release only male mosquitoes, in part because it is the females that bite humans and spread disease. Introducing large numbers of females could conceivably aid disease spread (Alphey et al., 2010) or contribute to nuisance biting. A challenge faced by the mass-rearing of mosquitoes in SIT programs is accurately separating males from females (sex separation). However, advances in the mechanical sorting of pupal and adult mosquitoes, as well as the availability of added safeguards such as irradiation, have seen renewed interest in controlling insect populations via the *Wolbachia*-IIT process (Atyame et al., 2011, Zheng et al., 2019). Whilst new methods can have a high degree of accuracy, occasional errors in such systems can lead to the release of *Wolbachia*-infected females. For IIT programs that rely on *Wolbachia* related CI effects, the release of female mosquitoes may have the unintended effect of population replacement rather than elimination. For a *Wolbachia* infection to become established within a population, it is generally thought that approximately 20% (depending upon the *Wolbachia* strain) of the population must become infected (Xi et al., 2005, Axford et al., 2016). Below this threshold, also known as the unstable equilibrium threshold (UET), the *Wolbachia*-infected genotype is likely to die out.

Historically, SIT and IIT control strategies have been underpinned by mathematical models, used to predict population dynamics of the target species. These models provide support tools for planning when, where and how many incompatible insects will be released into the wild population. The models of Knipling (Knipling, 1955, Knipling, 1959) provided the original mathematical frameworks on which successful insect control campaigns have been based. Knipling’s models used discrete-time dynamics and a simple model of geometric population growth, where the resulting suppression depends on an over-flooding ratio of sterile males to wild males (Knipling, 1955, Knipling, 1959). More recently, sophisticated SIT/IIT models have involved the modelling of genotypes and abundance via delay differential equations (Hancock et al., 2011), spatio-temporal advection-diffusion-reaction models with multiple life-stages (Dufourd and Dumont, 2013) and complex agent-based simulations (Magori et al., 2009). The majority of models used to study insect elimination processes via SIT/IIT are deterministic, ignoring the demographic randomness that is inherent to real populations.

Demographic stochasticity can be a significant source of uncertainty in populations, particularly, when populations are small and single births or deaths can have large effects on population dynamics (Hancock et al., 2019). Consequently, deterministic models do not quantify uncertain outcomes, such as the probability that a gene or *Wolbachia* symbiont will succeed or fail to enter a population. Models, such as that of Jansen et al. (2008) or Magori et al. (2009), are at a minimum stochastic but as we discuss below have certain attributes that affect their suitability for investigating important, general questions regarding the successful implementation of *Wolbachia*-IIT programs.

Simulations from stochastic models yield ensembles of population trajectories which can be used to estimate probabilities of particular events (e.g. whether population elimination will occur). Where, in addition to stochasticity in population dynamics, there is uncertainty in the initial conditions or parameters that govern the biological system, these quantities can be sampled from probability distributions which summarise the scientific literature or from prior expert knowledge (called *prior distributions*). Combining a stochastic population model with prior distributions allows us to estimate the probabilities of events, whilst acknowledging uncertainties in parameters and demographic stochasticity.

Markov chains are ubiquitous for studying population dynamics and such models have already been used to study the dynamics of *Wolbachia*-infection within a population. For example, Jansen et al. (2008) used a Moran process to estimate the probability of *Wolbachia* establishment after the release of a single infected female into a population of wild-type mosquitoes, but this process is not fit-for-purpose in SIT/IIT programs, since it does not allow for a declining population.

Markov Population Processes (MPPs; Kingman, 1969) are a class of continuous-time Markov chain models suitable for exploring the effects of *Wolbachia*-IIT population suppression, and therefore the risk of *Wolbachia* establishment. The Markov property dictates that in an MPP the times between events in the population follow an exponential distribution. Consequently, Markov population models typically assume that individual lifetimes are exponentially distributed. This is a reasonable assumption for mosquitoes in their natural habitat, where survival is frequently reported in terms of a constant daily mortality (Muir and Kay, 1998, Ritchie et al., 2013). For phenomena where lifetimes are not exponentially distributed, a sequence of states can be chained together so that the total time spent in those collective states is the sum of exponentially distributed random variables.

In this paper, we use an MPP model to address key questions related to the effective implementation of *Wolbachia*-IIT population suppression programs, namely:

i. What degree of fidelity in sex-separation is required to ensure that a *Wolbachia*-IIT population suppression program does not result in *Wolbachia* establishment and the replacement of the wild-type (non-infected) population?
ii. Does the release of a small number of females during a *Wolbachia*-IIT population suppression program imply a high probability of *Wolbachia* establishment?
iii. What types of release strategies are most effective for achieving population elimination without *Wolbachia* establishment?
iv. How does the establishment probability vary with the initial density of the *Wolbachia* infection in the population?

We recognise the important trade-off between increasing model complexity and a reduced ability to *defensibly* parameterize the model using what is known from the scientific literature so that it can be applied more generally. To overcome this, we present a novel MPP model that relies upon a relatively small set of parameters and is specifically designed to study *Wolbachia*-IIT population dynamics at the suburban block level. Importantly, our model is structured so that the parameters can be defensibly estimated from previous field studies (see Supplementary Material B, section B1).

## Methods

### Simulation of *Wolbachia*-IIT Programs

Our studies focus on simulating the repeated release of *Ae. aegypti*, infected with *w*AlbB *Wolbachia*, into a hypothetical population that is intended to represent a typical suburban block for Innisfail, northern Australia (−17.5226° S, 146.0285° E). Our intention is to obtain results that resemble a well-mixed biological population at the suburban block scale. Results obtained can be extrapolated to larger landscapes assuming independence between blocks. To allow for some heterogeneity between suburban block scale populations, our simulations of *Wolbachia*-IIT programs use adult equilibrium populations (prior to *w*AlbB releases) in the range 100 to 300 wild-type individuals/block. If we consider that a typical suburban block in Innisfail has about 20 houses, this represents between 5 and 15 adult *Ae. aegypti* mosquitoes per house, which is typical for North Queensland (Ritchie et al., 2013). For each simulated IIT program, we modelled one year of data with *w*AlbB releases occurring on Monday, Wednesday and Friday. After a year of *w*AlbB releases, we continued to model the population for a further year without releases, to monitor for any additional changes in population structure. After this two-year period, we classify each simulated trajectory as having an *IIT endpoint* of: (i) eliminated, (ii) established, (iii) indeterminate *Wolbachia* negative or (iv) indeterminate *Wolbachia* positive (see Table 1 for definitions). We assess the effectiveness of a *Wolbachia*-IIT program by calculating the proportions of simulations that reach these four IIT endpoints, with the elimination endpoint being the obvious goal.

**Table 1.**
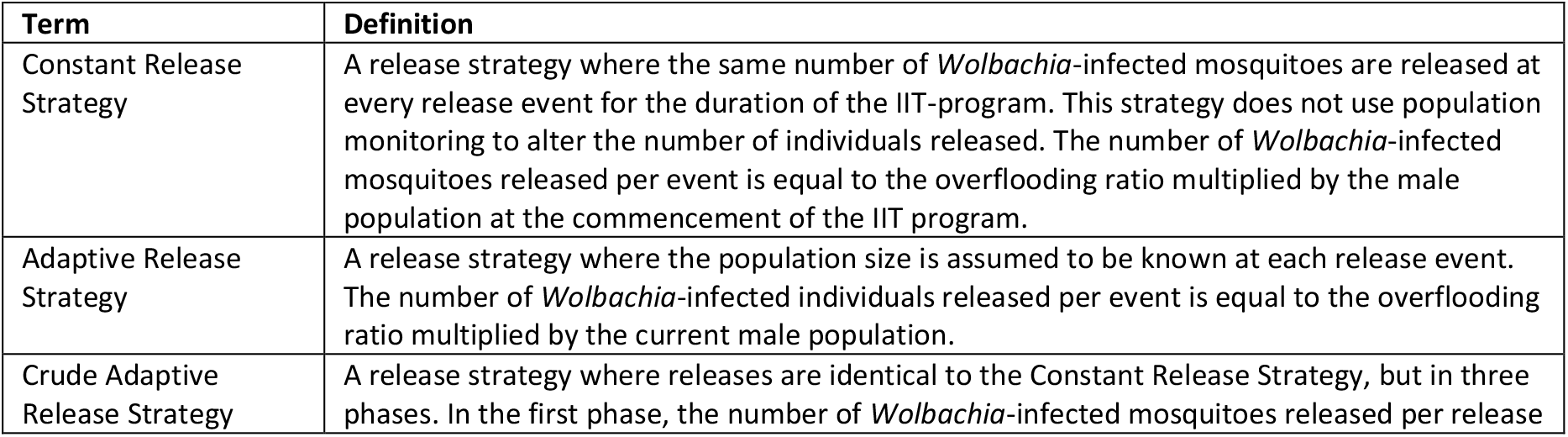
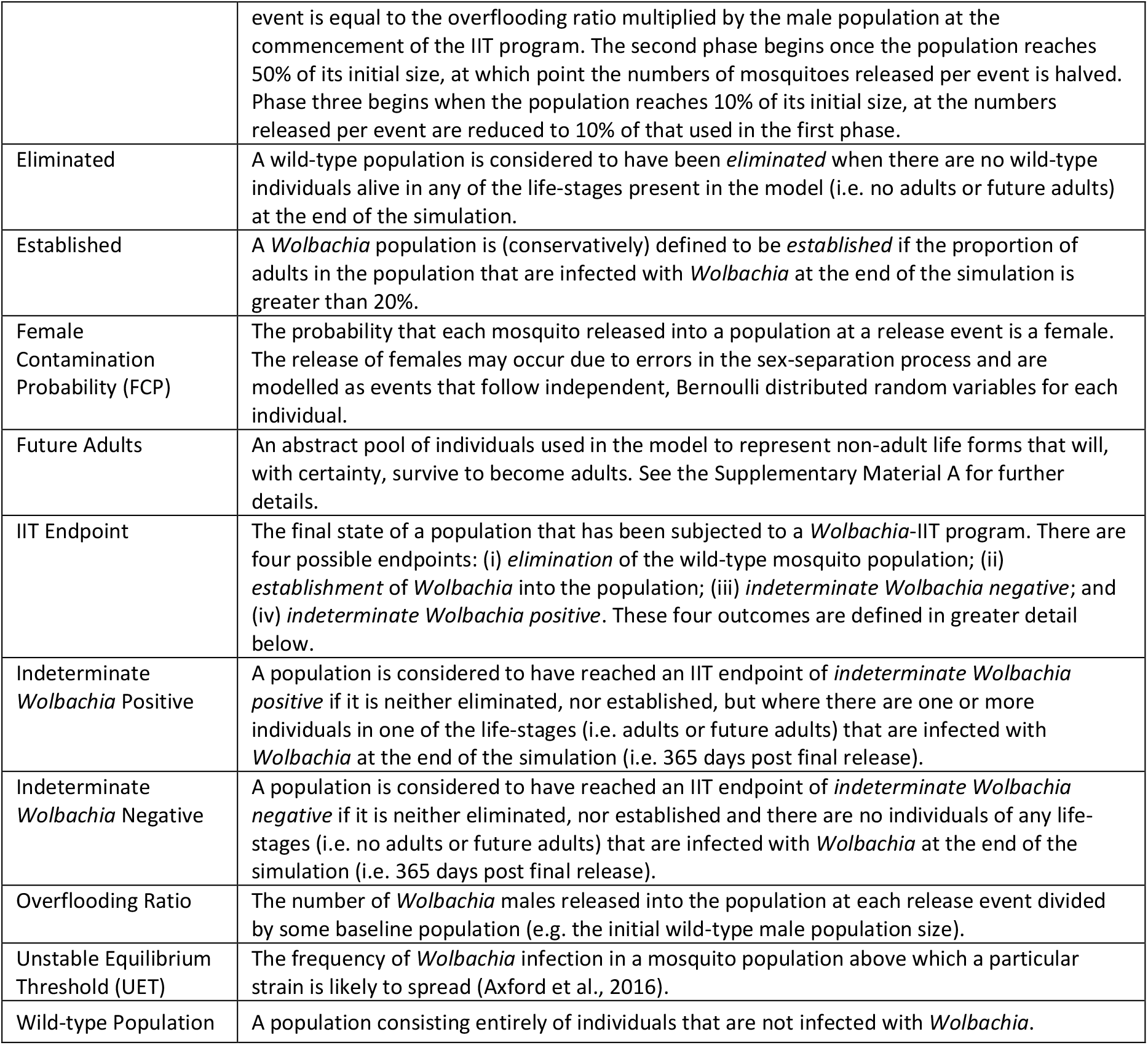
Glossary of modelling terminology.

We simulated *Wolbachia*-IIT programs using three release strategies (defined in Table 1): a constant release strategy; an adaptive release strategy; and a crude adaptive release strategy. For each of these release strategies we also considered two *overflooding ratios* (Table 1) of 5:1 and 15:1. These overflooding ratios scale the numbers of *w*AlbB individuals released under each of these three strategies. Release strategies and overflooding ratios in combination with four levels of sex-separation fidelity, determined by the *female contamination probability* (FCP; Table 1). We consider FCPs of 10^-4^, 10^-5^, 10^-6^ and 10^-7^, which span ranges that are theoretically achievable using next generation mechanical sex-separation approaches. We used 1000 simulations per scenario to assess the probability of *w*AlbB establishment and successful elimination. The additional simulations at the lowest FCP are to ensure that at least some population trajectories result in release of *w*AlbB females. Each set of simulations was used to study the probabilities of (i) unintended *w*AlbB establishment and (ii) wild-type elimination. Large numbers of simulations (15,000) per scenario were also used in conjunction with importance sampling (see Robert and Casella (2013)) to efficiently estimate establishment and elimination probabilities and their standard errors (see section B2 of Supplementary Material B).

In our simulations we do not assume that parameters are known precisely. Instead, we use a uniform probability distribution over plausible ranges for each of the parameters in our model (discussed in the subsequent section). For each simulation, a set of parameters is sampled from within these plausible ranges and checked to ensure that certain mandatory constraints are met (see section A4.4 of Supplementary Material A). In doing so, we ensure that our modelling results acknowledge parametric uncertainty and consider a plausible degree of variation that might be exhibited in a natural population. As such, simulations that result in a high probability of elimination can be considered robust to a range of parameterisations and population outcomes.

### The *Wolbachia*-IIT MPP Model

The states through which individuals in our model can progress are depicted in Figure 1. In total there are 2*k* + 8 different classes of individuals in the model and our MPP tracks the number of individuals in each class. For the wild-type and the *w*AlbB mosquito populations, there are: *k* classes of “future adults” (see Supplementary Material A) denoted *I*_*X*,1_, *I*_*X*,2_,…, *I_X,k_*, where *X* ∈ {*wild, w*Alb}; adult males denoted *M_X_*; unmated females denoted *F_X_*; and mated females denoted *F_X×Y_*, where *Y* ∈ {*wild, w*Alb}. The precise mathematical details of the MPP are provided in Supplementary Material A but we outline the general mechanics below. The model (as an installable R package) is publicly available at https://github.com/dpagendam/debugIIT.

**Figure 1.**
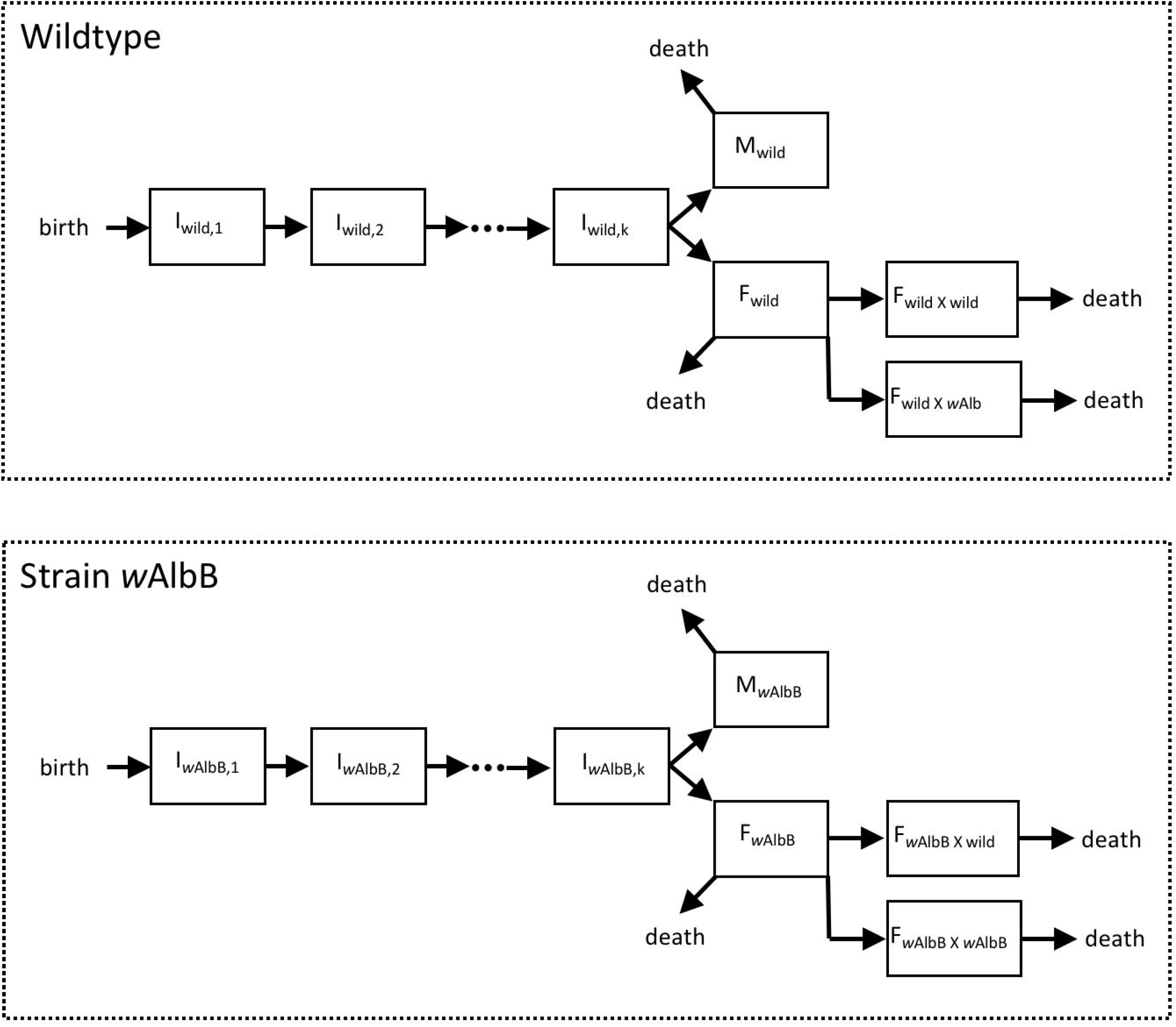
Model state transitions of wild-type and *w*AlbB-infected individuals from birth to death.

The modelling approach is designed to be parsimonious and is premised on the idea that, for a population to persist, we expect individuals to produce new adults at a rate that balances deaths in the population. To accommodate this, our model includes an abstract pool of “future adults” (see Table 1). That is, a cohort of individuals that are immature, but will survive to become new adults in the population. Mated females in the population give birth to new future adults and after a gamma-distributed, random period of time, each future adult emerges as a new adult that can mate and potentially give rise to new future adults. The rate at which mated females produce future adults is modelled to decrease as the pool of future adults increases in size and emulates density-dependent larval mortality without resorting to modelling the entire larval pool.

In Figure 1, we see that the birth of a new individual introduces it into the first class of the “future adults” (*I*_*X*,1_, individuals in each of the *k* future adult classes transition out of each state at rate *γ*. When an individual transitions out of state *I_X,k_*, it is allocated to the pool of males or unmated females with equal probability. The rate at which females are mated is assumed to be independent of the density of males in the population, and provided that there are males in the population, each female transitions to a mated state at rate *η*. We assume that females only mate once and that her mate is either wild-type with probability 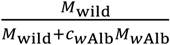, or *w*AlbB with probability 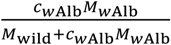. In other words, the rate at which females are mated by males is not density-dependent, but mate-type is. The parameter *c*_wAlb_ is the mating competitiveness coefficient or Fried’s index of *w*AlbB-infected individuals relative to wild-type individuals (Pagendam et al., 2018). We assume that CI between *w*AlbB males and wild-type females is 100% effective, so that only *F*_wild×wild_ females can give birth to wild-type future adults. Mated females of type *F*_*w*Alb×wild_ and *F*_*w*Alb×*w*Alb_ can give birth to *w*AlbB future adults. Each mated female adds new “future adults” to state *I*_*X*,1_ at rate 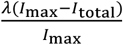, where 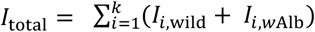 and *I*_max_ is a parameter that acts as a population ceiling on the number of future adults allowed within the population (see section A4 of Supplementary Material A). The parameter *λ* is the per capita rate at which mated females produce future adults in an empty niche (i.e. the intrinsic rate of population growth). This growth rate is modulated by a density-dependent term 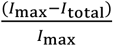 which can be thought of conceptually as a hypothetical proportion of niches for future adults which are currently unoccupied. Ultimately, this results in future adults being produced at a higher rate (respectively lower rate) when the density of future adults is low (respectively high). In essence, this provides a simple representation of higher larval mortality at higher larval population densities (Dye, 1984, Hancock et al., 2016). Section A4.2 of Supplementary Material A outlines how the abstract parameter, *I*_max_, can be identified from a small number of easily estimated quantities presented in Table B1.1 of Supplementary Material B.

Our decision to model the female mating rate as a density-independent process stems from the assumption that adult males and females have little difficulty finding each other at the level of a suburban block and that the mating rate is not entirely governed by random interactions and may involve behavioural bottlenecks. Therefore, we argue that the rate of female mate selection is not simply increased monotonically by the presence of more males. There is also good scientific evidence to suggest that adult males and females can find each other by means of sensory cues (Johnson and Ritchie, 2015), so that the female mating rate does not necessarily diminish greatly at low adult male densities. In modelling the probability of *w*AlbB establishment we do introduce a conservative assumption that any *w*AlbB females accidentally released will have been premated with a *w*AlbB male. This reflects an assumption of prolonged exposure of each *w*AlbB female to many *w*AlbB males in isolation.

We note that the mathematical model presented in Supplementary Material A is more general than that presented here and accommodates populations with multiple strains of *Wolbachia* infection. Sections A2 and A4 of Supplementary Material A also describe how our model relies on a set of primary and secondary parameters. Primary parameters are quantities that the user must specify to run the model, whilst secondary parameters are not defined by the user but are uniquely defined by the values of the primary parameters and the equilibrium dynamics of the model. Parameter ranges used as uniform prior probability distributions for simulations are provided in Table B1.1 (Supplementary Material B).

Most of the parameters listed in Table B1.1 (Supplementary Material B) can be estimated from field or laboratory data and we provide references that were used for these purposes. Some of these parameters require some additional detail, namely *k* and *γ*, which together dictate that the mean time between a female laying an egg containing a future adult and the emergence of that new adult can range between 10 and 40 days (consistent with Hancock et al. (2016)) with the distribution for the time spent in immature form following a gamma distribution. In addition, *p*_mated_ was estimated in a series of (unpublished) experiments conducted over five different weeks using the Rhodamine B marking approach of Johnson et al. (2017), where wild-type females were captured and the spermatheca dissected to ascertain whether mating had taken place.

### Unstable Equilibrium Threshold (UET) and the Probability of *w*AlbB Establishment

To ensure that our model was biologically defensible, we checked for the existence of a UET of the type discussed by Hoffmann et al. (2011) and demonstrated in laboratory populations by Axford et al. (2016). These studies suggest that there should be a high probability of *w*AlbB becoming established when the number of *w*AlbB infected adults introduced exceeds a *w*AlbB frequency of approximately 20%. There have been a number of theoretical investigations employing deterministic models to study the UET (Turelli and Hoffmann, 1991, Turelli, 1994, Barton and Turelli, 2011). To examine the existence of a UET in our stochastic system, we ran simulations of our model using a wild-type population with an equilibrium of 400 adult individuals (to maintain similarity to the cage experiments of Axford et al. (2016)). We generated simulations of our population where *w*AlbB adults in a 1:1 sex ratio were introduced to this population, representing between 5% to 40% (in intervals of 5%) of the total adult population. For each proportion of introduced *w*AlbB adults, we simulated 100 population trajectories over a five year period. Simulations were conducted under two further scenarios: the first where none of the released *w*AlbB females had been mated prior to release, and the second where all *w*AlbB females were assumed to have been mated by *w*AlbB males prior to release. The establishment probabilities for each set of 100 simulations were estimated as the proportions of trajectories where the wild-type populations were completely replaced by the *w*AlbB-infected strain. Confidence intervals around the estimated establishment probabilities were derived using profile likelihood (i.e. by inverting the likelihood ratio test statistic).

## Results

We ran a total of 114,000 individual simulations each taking an average of 17 minutes to run on a single CPU core, with 90,000 of those simulations for importance sampling estimates of establishment and eradication probabilities. This amounted to a total of 1,345 days of HPC compute time with runs parallelised to perform up to 5,000 concurrent simulations at maximum capacity. Each simulation was allocated 8 GB of RAM and was run on an Intel Xeon E5-2670 V3 processor running at 2.6Ghz.

In addressing the four questions proposed by this study, we focused on a small set of plots and tables that demonstrate the following key results:

i. To ensure a very low probability (< 0.01) of *w*AlbB establishment on a single suburban block, an FCP of 10^-6^ or less is advisable, but this probability is also a function of the number of mosquitoes released and is affected by the release strategy and overflooding ratio (Figure 2).
ii. The release of a small number of females over the duration of a *Wolbachia*-IIT program, does not necessarily imply that there is a high probability of establishment (Figure 3).
iii. To ensure a very low probability (< 0.01) of any block-level *w*AlbB establishment over an area spanning as many as 100 suburban blocks, it is advisable to adopt the crude adaptive release strategy with an FCP of 10^-7^ or less using a 5:1 overflooding ratio (Figure B3.1 and Table B2.1 of the Supplementary Material B).
iv. Of the three release strategies examined, the best IIT outcomes (i.e. high probability of eradiation and low probability of establishment) were achieved using a crude adaptive release strategy with a 5:1 overflooding ratio at lower FCPs (Figure 2, Tables B2.1 and B2.2 of Supplementary Material B).
v. The stochastic population model developed and used in this study demonstrates a similar (UET) to that observed previously in laboratory cage experiments (Figure 4 and Figure B3.2 of Supplementary Material B).

**Figure 2.**
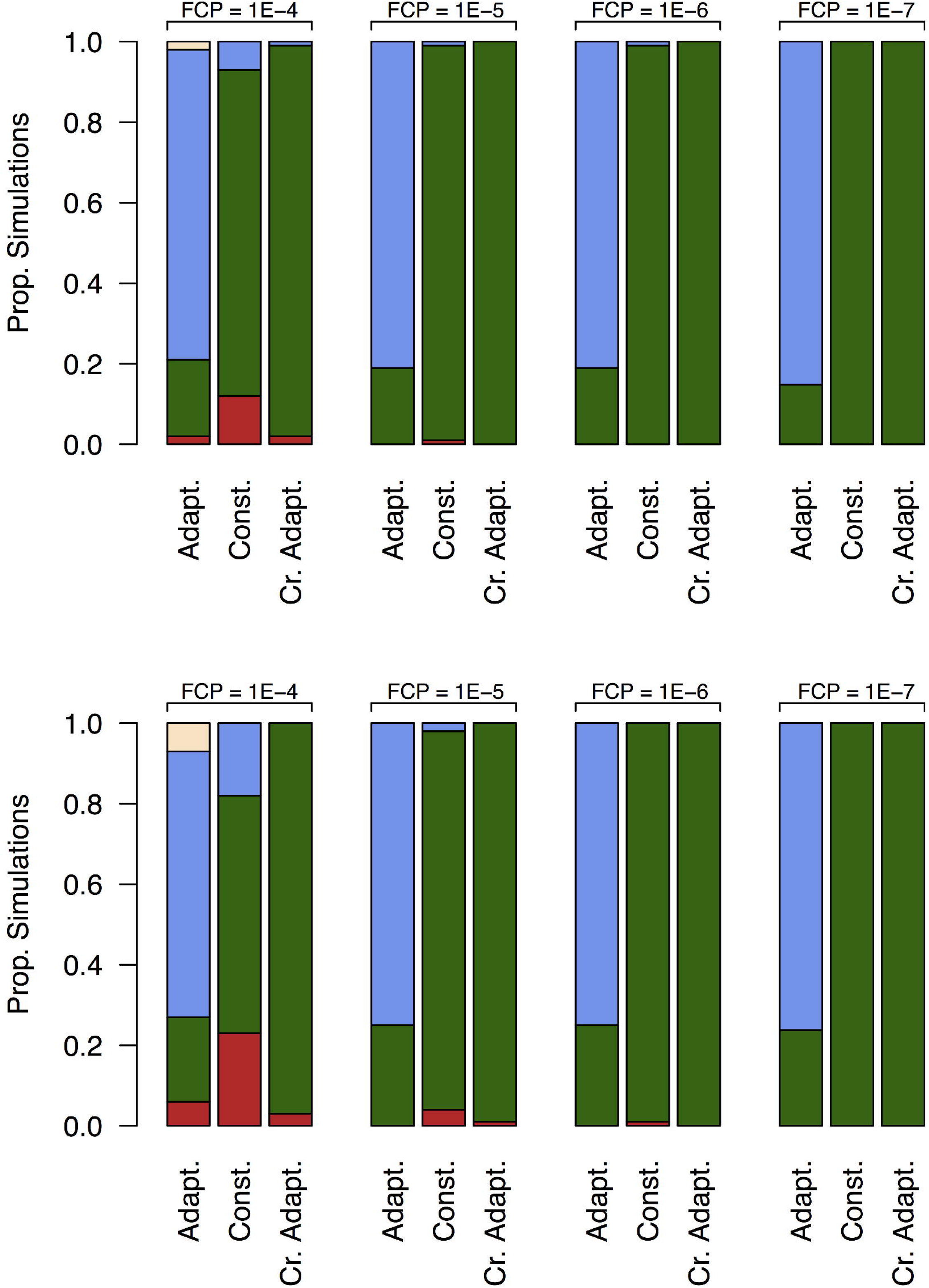
Proportions of simulations that either resulted in: *w*AlbB establishment (red), successful wild-type elimination (green); indeterminate *w*AlbB negative (blue); and indeterminate *w*AlbB positive (yellow) for scenarios based on different FCPs and release strategies at 5:1 (top row) and 15:1 (bottow row) overflooding. Results suggest that a lower FCP leads to higher elimination rates. At the lower FCP, the best results appear to be for the constant and crude adaptive release strategies.

**Figure 3.**
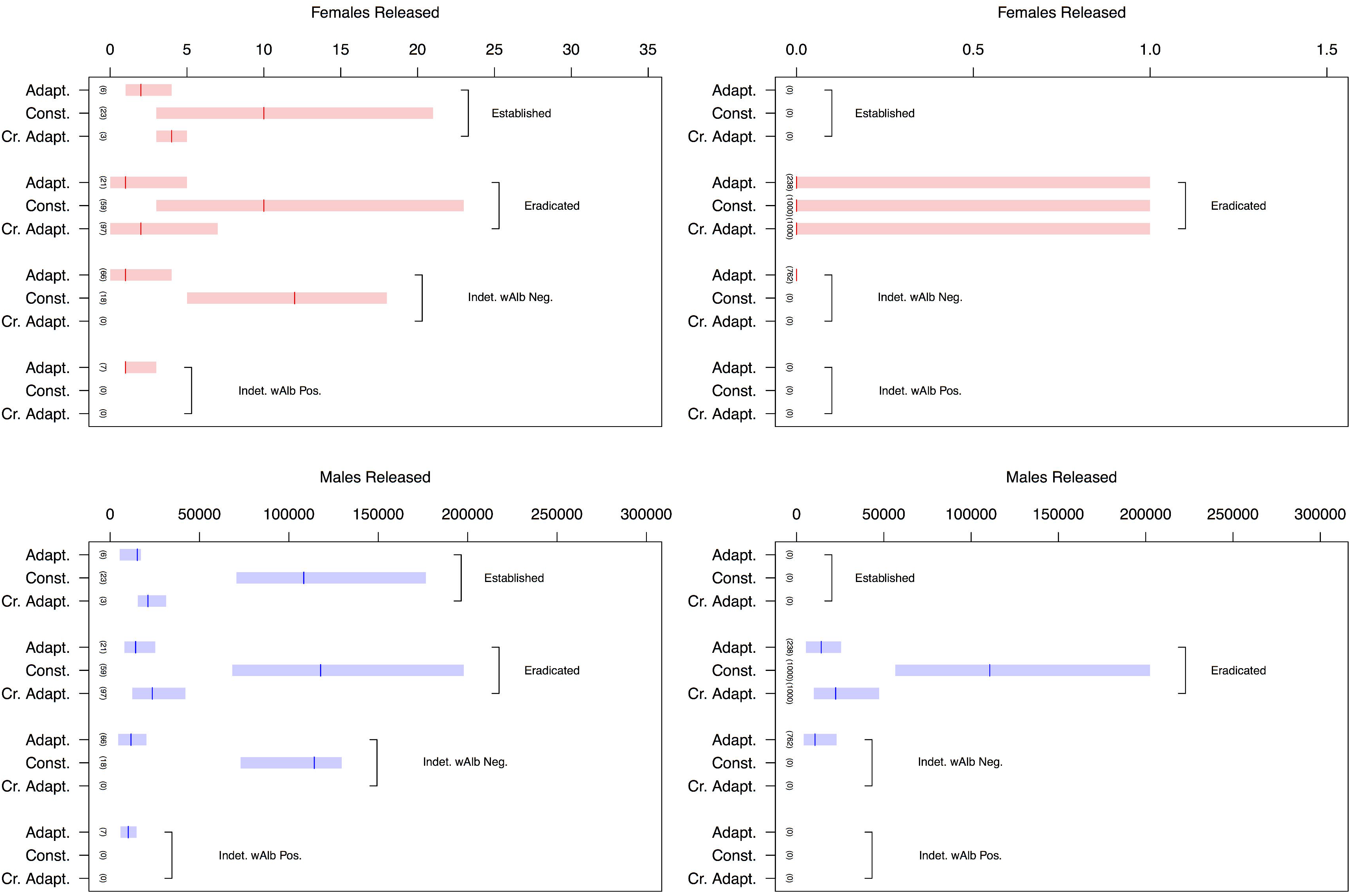
Statistics for the number of males and females released using a 15:1 overflooding and FCP of 10^-7^ (top row); and 10^-4^ (bottom row). Rectangles span the minima and maxima for each set of simulations; horizontal red and blue lines show the medians for each set of simulations; and numbers in brackets show the number of simulations (out of 1000) that resulted in each IIT endpoint.

**Figure 4.**
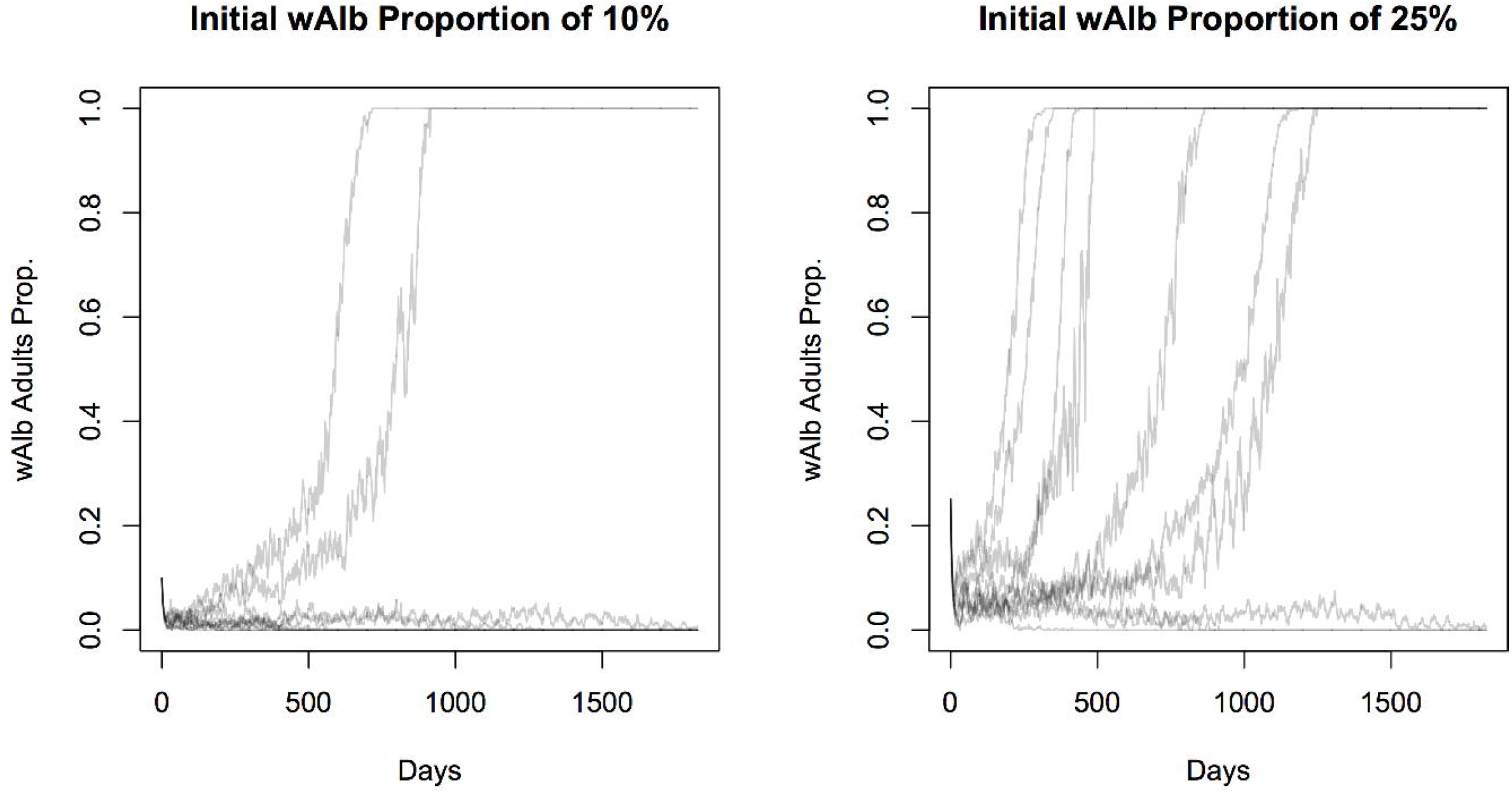
A random sample of ten simulated trajectories of the cage experiment scenario (no *w*AlbB prerelease mating) at *w*AlbB initial proportions of 10% (left) and 25% (right).

The proportions of simulations whose IIT endpoints were classed as either successful elimination, *w*AlbB establishment, indeterminate *w*AlbB negative, or indeterminate *w*AlbB positive, are shown in Figure 2. These proportions are plotted for different sex separation fidelities, release strategies and target populations. We observed a clear decline in the incidence of establishment events with lower FCPs. Furthermore, the adaptive release strategy demonstrated a high likelihood of achieving an endpoint of indeterminate *w*AlbB negative across all FCPs and overflooding ratios studied. In addition to Figure 2, Tables B2.1 and B2.2 of Supplementary Material B provide importance sampling estimates of the *w*AlbB establishment probabilities. Notably, this statistical approach provides statistically efficient estimates of the establishment probability, even when simulated establishment events are rarely encountered at the FCP of 10^-7^.

To address the question of whether the accidental release of a small number of *w*AlbB females is likely to result in a high probability of establishment, we classified simulations by their IIT endpoints. Figure 3 dissects some of these simulated trajectories (15:1 overflooding, with FCP of 10^-6^ and 10^-7^) in this way. The plots serve to provide useful statistics (minimum, median and maximum) for the numbers of males and females released for each of the four *Wolbachia*-IIT endpoints.

Finally, we demonstrated that our model (and its parameterisation) yields simulated population dynamics consistent with observations of unstable equilibria in biological systems. In a population having a stable wildtype equilibrium of 400 individuals (i.e. *K_wild_* = 400) we simulated trajectories with different initial proportions of *w*AlbB and wild-type individuals. We display the first ten simulated population trajectories summarised as the proportion of adults infected with *w*AlbB for both frequency scenarios where no pre-release mating has occurred (Figure 4). In those simulations where there was an initial 25% *w*AlbB frequency, 62 out of 100 simulations ended in *w*AlbB establishment, whereas at the smaller initial frequency of 10% *w*AlbB, only 25 out of 100 trajectories established. The point at which the establishment probability exceeded a 50% probability occurred at between 15 and 20% *w*AlbB for no pre-release mating and between 10 and 15% where pre-release mating was present. To quantify the differences in *w*AlbB establishment probabilities between mated and unmated introduced females, Figure B3.2 of Supplementary Material B displays the estimated establishment probabilities for each *w*AlbB frequency level using 100 simulated populations at each of these levels.

## Discussion

The development of the MPP presented here has allowed a deeper understanding of the uncertainties associated with female contamination frequencies in *Wolbachia*-IIT programs and highlights the benefits of adopting stochastic rather than deterministic modelling frameworks. Importantly, our model highlights the attributes of *Wolbachia*-IIT programs that yield high probabilities of eliminating a population without *Wolbachia* establishment: (i) very high-fidelity sex separation and (ii) overflooding ratios and release strategies that employ only moderate numbers of *Wolbachia*-infected males. The former point highlights a need for technologies that can effectively discriminate between sexes and sort mosquitoes without reducing their fitness. The latter highlights a need to understand the long-term outlook when conducting such IIT programs. It follows that releasing an excess of *Wolbachia*-infected male mosquitoes will undermine a program by increasing the potential number of females accidentally released, while increasing the demand on rearing facilities. This is particularly the case as the first area-wide SIT/IIT programs have typically employed a constant or increasing release rate over time (Kittayapong et al., 2019, Crawford et al., 2020). If the rate at which females are mated is density-independent and only the mate type is density-dependent (as we have assumed in our model), then very high overflooding ratios hold little strategic value over moderate ones. Figure 3 shows that releasing very large numbers of mosquitoes (e.g. 15:1 overflooding under the Constant Release Strategy compared to the Crude Adaptive Release Strategy) tends to result in a greater likelihood of *w*AlbB establishment.

Of the three release strategies and two overflooding ratios simulated, the best performing approach was the Crude Adaptive Release Strategy in conjunction with a 5:1 overflooding ratio. The Adaptive Release Strategy appeared to be ineffective at both the 5:1 and 15:1 overflooding ratios, as the number of males released over one year resulted in many outcomes of “indeterminate *Wolbachia* negative” (population suppression without elimination or *Wolbachia* establishment). It is possible that this approach may achieve satisfactory elimination when releases are scheduled over a longer period of time, however, given the observed success of the Crude Adaptive Release Strategy at 5:1 overflooding, we did not pursue this further. Furthermore, implementing the Adaptive Release Strategy seems difficult in practice, requiring accurate, real-time surveillance and population size estimates to tailor release numbers appropriately. The Constant Release Strategy also showed encouraging results at the lowest FCP and 5:1 overflooding ratio. However, we estimate the establishment probability for the Crude Adaptive Release Strategy to be order of magnitude lower than for constant overflooding and is therefore preferable for mitigating the risk of *Wolbachia* establishment (particularly when applied over larger urban areas). An aspect that was not investigated in our modelling is the probability of establishment when a female is released towards the end of an IIT program, when compared to earlier stages. This reflects a reduced barrier to entering a niche when larval densities are lower because of reduced density-dependent mortality.

The simulations in this study were intended to represent suburban blocks in north Queensland, Australia, which typically consist of ~20 houses. Since a population suppression program would typically involve treating larger urban areas, the probability of one or more blocks resulting in *w*AlbB establishment will increase over regions consisting of multiple blocks. Under a simplifying assumption that all blocks in a treated region are independent, non-interacting subpopulations, the probability of *w*AlbB establishing on one or more blocks is calculated as *π* = 1 – (1 – *p*)^*n*^, where *p* is the block level establishment probability, *n* is the number of blocks treated with *w*AlbB. For a hypothetical region consisting of 100 blocks, a block-level *w*AlbB establishment rate of less than 10^-4^ would be required in order to ensure that the *overall* probability of one or more blocks having an establishment event was less than 0.01. Based on this, the most suitable release strategy investigated would be a Crude Adaptive Release Strategy with an FCP of 10^-7^ and an overflooding ratio of 5:1 or 15:1, which have block-level *w*AlbB establishment rates of 1.546 x 10^-5^ and 4.159 x 10^-5^ respectively.

We believe that our estimates of the probability of establishment risk are conservative, since we assumed that all *w*AlbB females were mated and yielded viable offspring. Through these simulations we hope to debunk the legitimate concern amongst biologists that IIT programs may not be robust to the release of a small number of *Wolbachia*-infected females and the subsequent risk of establishment. The probabilistic evidence that we have presented here suggests that *Wolbachia* establishment depends on many factors, including the population size at the time of accidental female release, the release strategy, and the overflooding ratio employed. In many of our simulated IIT programs, where females were accidentally released, *Wolbachia* failed to establish, primarily due to the presence of the UET coupled with demographic stochasticity. To demonstrate this point, Figure 3 shows that as many as 23 females were released in one of the trajectories at the 10^-4^ FCP, and yet elimination was achieved, and the population was successfully eradicated. Major factors affecting establishment in the population are: whether a female survives long enough post-release to produce new individuals; and the population density of larvae that reduces the probability of an individual female replacing herself. As such, it is important to acknowledge that if a small number of *w*AlbB females were detected in the population then the chance of establishment would be further decreased if releases ceased for a period.

To increase confidence in the population dynamics exhibited in our MPP model, we endeavoured to replicate a scenario similar to the cage experiments of Axford et al. (2016). The key findings from our simulations were: (i) for pre-mated *w*AlbB individuals, the majority of simulations resulted in establishment where the initial *w*AlbB frequency was above 20% and (ii) for *w*AlbB individuals without pre-mating, the majority of simulations established above an initial 15% *w*AlbB frequency. These results show a general agreement with the Axford et al. (Axford et al., 2016) laboratory cage experiments where a high probability (five out of five laboratory populations) of *w*AlbB establishment occurred when the initial frequency of *w*AlbB was 22%. Likewise, both this study and Axford et al. (2016) show a much lower probability of establishment (one out of five populations in the latter study) when the initial frequency of *w*AlbB in the population was 10%.

There are clear benefits to using stochastic models over deterministic models in population biology for addressing questions related to probability and risk. However, in addressing our questions about *w*AlbB establishment risk, we required a model that was fit for purpose and easily parameterised. Existing stochastic models were deemed unsuitable. For example, the model of Jansen et al. (2008) assumes that the population size remains constant over time (which is violated during IIT programs) and that populations evolve in discrete-time through non-overlapping generations; and Magori et al. (2009) requires the identification of many spatially-explicit parameters (e.g. numbers and locations of breeding containers). The simple MPP model introduced in this work relied on a simple set of parameters that could be determined from the literature and past field data and was general enough to draw conclusions about a typical suburban block in Innisfail, Australia. We believe that this study demonstrates the usefulness of such MPP models for making important *in silico* observations about what factors might be important in designing an effective SIT/IIT strategy. Furthermore, our model accounts for lags between a female giving birth to a new individual and that individual’s eventual emergence as an adult, avoiding the complicated processes of tracking every individual in each life-stage of a mosquito population or conditioning our results on detailed spatial information (e.g. location and number of containers) used in other mosquito modelling system such as CimSim (Focks et al., 1993a, Focks et al., 1993b) and SkeeterBuster (Magori et al., 2009). Importantly, we avoid explicitly modelling all life stages because: (i) the life-history parameters of juveniles appear to be less well understood in the field and (ii) this would drastically increase the numbers of individuals that would need to be stochastically modelled in the population resulting in significant computational overheads. Furthermore, modelling every individual during each life-cycle stage in a population may be of little real benefit since the vast majority of juveniles do not reach adulthood due to density-dependent larval mortality (Dye, 1984, Hancock et al., 2016). We believe that the parsimonious approach adopted in our MPP model simplifies these processes and can be conceptually and computationally advantageous when running simulations over much larger spatial areas.

## Conclusion

From the perspective of designing an effective IIT program, our study suggests that to achieve elimination while avoiding establishment at large spatial scales (i.e. many suburban blocks), it is advisable to have a high-fidelity sex separation method with an FCP of 10^-7^ or less. Our results favour the crude adaptive release strategy at a 5:1 overflooding ratio to achieve both a low establishment probability and high elimination probability at the 10^-7^ FCP level. Our simulations indicate that releasing a relatively small number of *w*AlbB females during a program does not automatically render it a failure.

## Data Accessibility

The model code is publicly available as an installable R package at https://github.com/dpagendam/debugIIT.

## References

Alphey, L., Benedict, M., Bellini, R., Clark, G. G., Dame, D. A., Service, M. W. & Dobson, S. L. 2010. Sterile-Insect Methods for Control of Mosquito-Borne Diseases: An Analysis. Vector-Borne and Zoonotic Diseases, 10, 295–311.

Atyame, C. M., Pasteur, N., Dumas, E., Tortosa, P., Tantely, M. L., Pocquet, N., Licciardi, S., Bheecarry, A., Zumbo, B., Weill, M. & Duron, O. 2011. Cytoplasmic Incompatibility as a Means of Controlling Culex pipiens quinquefasciatus Mosquito in the Islands of the SouthWestern Indian Ocean. PLOS Neglected Tropical Diseases, 5, e1440.

Axford, J. K., Ross, P. A., Yeap, H. L., Callahan, A. G. & Hoffmann, A. A. 2016. Fitness of wAlbB Wolbachia Infection in Aedes aegypti: Parameter Estimates in an Outcrossed Background and Potential for Population Invasion. The American Journal of Tropical Medicine and Hygiene, 94, 507–516.

Barton, N. H. & Turelli, M. 2011. Spatial Waves of Advance with Bistable Dynamics: Cytoplasmic and Genetic Analogues of Allee Effects. The American Naturalist, 178, E48–E75.

Crawford, J. E., Clarke, D. W., Criswell, V., Desnoyer, M., Cornel, D., Deegan, B., Gong, K., Hopkins, K. C., Howell, P., Hyde, J. S., Livni, J., Behling, C., Benza, R., Chen, W., Dobson, K. L., Eldershaw, C., Greeley, D., Han, Y., Hughes, B., Kakani, E., Karbowski, J., Kitchell, A., Lee, E., Lin, T., Liu, J., Lozano, M., Macdonald, W., Mains, J. W., Metlitz, M., Mitchell, S. N., Moore, D., Ohm, J. R., Parkes, K., Porshnikoff, A., Robuck, C., Sheridan, M., Sobecki, R., Smith, P., Stevenson, J., Sullivan, J., Wasson, B., Weakley, A. M., Wilhelm, M., Won, J., Yasunaga, A., Chan, W. C., Holeman, J., Snoad, N., Upson, L., Zha, T., Dobson, S. L., Mulligan, F. S., Massaro, P. & White, B. J. 2020. Efficient production of male Wolbachia-infected Aedes aegypti mosquitoes enables large-scale suppression of wild populations. Nature Biotechnology, 38, 482–492.

Dufourd, C. & Dumont, Y. 2013. Impact of environmental factors on mosquito dispersal in the prospect of sterile insect technique control. Computers & Mathematics with Applications, 66, 1695–1715.

Dyck, V. A., Hendrichs, J. & Robinson, A. S. 2006. Sterile insect technique: principles and practice in area-wide integrated pest management, Springer.

Dye, C. 1984. Competition amongst larval Aedes aegypti: the role of interference. Ecological Entomology, 9, 355–357.

Focks, D. A., Haile, D. G., Daniels, E. & Mount, G. A. 1993a. Dynamic Life Table Model for Aedes aegypti (Diptera: Culicidae): Analysis of the Literature and Model Development. Journal of Medical Entomology, 30, 1003–1017.

Focks, D. A., Haile, D. G., Daniels, E. & Mount, G. A. 1993b. Dynamic Life Table Model for Aedes aegypti (Diptera: Culicidae): Simulation Results and Validation. Journal of Medical Entomology, 30, 1018–1028.

Gilles, J. R. L., Schetelig, M. F., Scolari, F., Marec, F., Capurro, M. L., Franz, G. & Bourtzis, K. 2014. Towards mosquito sterile insect technique programmes: Exploring genetic, molecular, mechanical and behavioural methods of sex separation in mosquitoes. Acta Tropica, 132, S178–S187.

Hancock, P. A., Ritchie, S. A., Koenraadt, C. J. M., Scott, T. W., Hoffmann, A. A. & Godfray, H. C. J. 2019. Predicting the spatial dynamics of Wolbachia infections in Aedes aegypti arbovirus vector populations in heterogeneous landscapes. 56, 1674–1686.

Hancock, P. A., Sinkins, S. P. & Godfray, H. C. J. 2011. Population Dynamic Models of the Spread of Wolbachia. The American Naturalist, 177, 323–333.

Hancock, P. A., White, V. L., Callahan, A. G., Godfray, C. H. J., Hoffmann, A. A. & Ritchie, S. A. 2016. Density-dependent population dynamics in Aedes aegypti slow the spread of wMel Wolbachia. Journal of Applied Ecology, 53, 785–793.

Hendrichs, J., Ortiz, G., Liedo, P. & Schwarz, A. Six years of successful medfly program in Mexico and Guatemala. Fruit flies of economic importance. Proc. CEC/IOBC Int. Symp., November 1982, 1983. 353–365.

Hoffmann, A. A., Montgomery, B. L., Popovici, J., Iturbe-Ormaetxe, I., Johnson, P. H., Muzzi, F., Greenfield, M., Durkan, M., Leong, Y. S., Dong, Y., Cook, H., Axford, J., Callahan, A. G., Kenny, N., Omodei, C., Mcgraw, E. A., Ryan, P. A., Ritchie, S. A., Turelli, M. & O’NEILL, S. L. 2011. Successful establishment of Wolbachia in Aedes populations to suppress dengue transmission. Nature, 476, 454–457.

Jansen, V. A. A., Turelli, M. & Godfray, H. C. J. 2008. Stochastic spread of Wolbachia. Proceedings of the Royal Society B: Biological Sciences, 275, 2769–2776.

Johnson, B. J., Mitchell, S. N., Paton, C. J., Stevenson, J., Staunton, K. M., SNoad, N., Beebe, N., White, B. J. & Ritchie, S. A. 2017. Use of rhodamine B to mark the body and seminal fluid of male Aedes aegypti for mark-release-recapture experiments and estimating efficacy of sterile male releases. PLOS Neglected Tropical Diseases, 11, e0005902.

Johnson, B. J. & Ritchie, S. A. 2015. The Siren’s Song: Exploitation of Female Flight Tones to Passively Capture Male Aedes aegypti (Diptera: Culicidae). Journal of Medical Entomology, 53, 245–248.

Kingman, J. F. C. 1969. Markov population processes. Journal of Applied Probability, 6, 1–18.

Kittayapong, P., Kaeothaisong, N.-O., Ninphanomchai, S. & Limohpasmanee, W. 2018. Combined sterile insect technique and incompatible insect technique: sex separation and quality of sterile Aedes aegypti male mosquitoes released in a pilot population suppression trial in Thailand. Parasites & Vectors, 11, 657.

Kittayapong, P., Ninphanomchai, S., Limohpasmanee, W., Chansang, C., Chansang, U. & Mongkalangoon, P. 2019. Combined sterile insect technique and incompatible insect technique: The first proof-of-concept to suppress Aedes aegypti vector populations in semi-rural settings in Thailand. PLoS neglected tropical diseases, 13, e0007771–e0007771.

Knipling, E. F. 1955. Possibilities of Insect Control or Eradication Through the Use of Sexually Sterile Males. Journal of Economic Entomology, 48, 459–462.

Knipling, E. F. 1959. Sterile-Male Method of Population Control. Successful with some insects, the method may also be effective when applied to other noxious animals, 130, 902–904.

Laven, H. 1967. Eradication of Culex pipiens fatigans through Cytoplasmic Incompatibility. Nature, 216, 383–384.

Magori, K., Legros, M., Puente, M. E., Focks, D. A., Scott, T. W., Lloyd, A. L. & Gould, F. 2009. Skeeter Buster: A Stochastic, Spatially Explicit Modeling Tool for Studying Aedes aegypti Population Replacement and Population Suppression Strategies. PLOS Neglected Tropical Diseases, 3, e508.

Muir, L. E. & Kay, B. H. 1998. Aedes aegypti survival and dispersal estimated by mark-release-recapture in northern Australia. The American Journal of Tropical Medicine and Hygiene, 58, 277–282.

Pagendam, D., Snoad, N., Yang, W.-H., Segoli, M., Ritchie, S., Trewin, B. & Beebe, N. 2018. Improving Estimates of Fried’s Index from Mating Competitiveness Experiments. Journal of Agricultural, Biological and Environmental Statistics, 23, 446–462.

Ritchie, S. A., Montgomery, B. L. & Hoffmann, A. A. 2013. Novel Estimates of Aedes aegypti (Diptera: Culicidae) Population Size and Adult Survival Based on Wolbachia Releases. Journal of Medical Entomology, 50, 624–631.

Robert, C. & Casella, G. 2013. Monte Carlo statistical methods, Springer Science & Business Media.

Turelli, M. 1994. Evolution of incompatibility-inducing microbes and their hosts. Evolution, 48, 1500–1513.

Turelli, M. & Hoffmann, A. A. 1991. Rapid spread of an inherited incompatibility factor in California Drosophila. Nature, 353, 440–442.

Vargas-Terán, M., Hofmann, H. C. & Tweddle, N. E. 2005. Impact of Screwworm Eradication Programmes Using the Sterile Insect Technique. In: Dyck, V. A., Hendrichs, J. & Robinson, A. S. (eds.) Sterile Insect Technique: Principles and Practice in Area-Wide Integrated Pest Management. Dordrecht: Springer Netherlands.

Vreysen, M. J., Saleh, K. M., Ali, M. Y., Abdulla, A. M., Zhu, Z. R., Juma, K. G., Dyck, V. A., Msangi, A. R., Mkonyi, P. A. & Feldmann, H. U. 2000. Glossina austeni (Diptera: Glossinidae) eradicated on the island of Unguja, Zanzibar, using the sterile insect technique. Journal of economic entomology, 93, 123–135.

Xi, Z., Khoo, C. C. H. & Dobson, S. L. 2005. *Wolbachia* establishment and invasion in an *Aedes aegypti* laboratory population. Science, 310, 326–328.

Zheng, X., Zhang, D., Li, Y., Yang, C., Wu, Y., Liang, X., Liang, Y., Pan, X., Hu, L., Sun, Q., Wang, X., Wei, Y., Zhu, J., Qian, W., Yan, Z., Parker, A. G., Gilles, J. R. L., Bourtzis, K., Bouyer, J., Tang, M., Zheng, B., Yu, J., Liu, J., Zhuang, J., Hu, Z., Zhang, M., Gong, J.-T., Hong, X.-Y., Zhang, Z., Lin, L., Liu, Q., Hu, Z., Wu, Z., Baton, L. A., Hoffmann, A. A. & Xi, Z. 2019. Incompatible and sterile insect techniques combined eliminate mosquitoes. Nature, 572, 56–61.

