## Supplementary A for "Modelling the *Wolbachia* Incompatible Insect Technique: strategies for effective mosquito population elimination"

### *A1. Background Theory: Markov Population Process*

We use a stochastic model, known as a Markov population process (MPP; see Kingman, 1969) to model the biological system and the dynamics of *Wolbachia*-induced IIT programs. MPPs are a type of continuous time Markov chain (CTMC) with a multidimensional state vector that takes discrete (integer) values. The state vector $Y_t$ of the MPP, is a random vector, modelling the numbers of individuals in a population at time $t$, where the population consists of collection of classes (e.g. life stages or organism types).

The MPP is governed by a collection of fixed rates which are gathered into a matrix, $Q = (q(s, s^*), s, s^* \in S)$ and where $q(s, s^*)$ represents the rate of transition from state $s$ to state $s^*$ for $s \neq s^*$. The diagonal elements of $Q$ are defined as $q(s, s) = -q(s)$, where $q(s) = -\sum_{s^* \neq s} q(s, s^*)$, is the total rate at which we leave state $s$; $S$ is the state space and represents all possible configurations (states) of the process ($S$ being some subset of $\mathbb{Z}^D$, where D is the number of elements in $Y_t$). In the context of the model presented herein, the state space is finite ($|S| < \infty$) and the dynamics of the state probability vector $p$ evolve according to $dp/dt = pQ$, where the elements of $p$ are the probabilities that the system is found in a particular state at a point in time. In the current model, and many other ecological models with finite state spaces, the transition function $P(t)$ of the system is obtained as $P(t) = \exp(Qt)$ (where exp( ) is the matrix exponential). $P(t)$ is a matrix with entries $p_{ij}$ that give the probability of moving from state $i$ to state $j$ over the time interval of length $t$.

When the state space, $S$, is very large, it may be inefficient to work with the matrices $Q$ and $P$ for investigating certain research questions. In some cases, sampling trajectories from the system defined by the transition rates in $Q$ may provide useful insights. Fortunately, simulation from these systems is straight-forward and the algorithm for doing so is referred to as the Doob-Gillespie Algorithm (Doob, 1945; Gillespie, 1976). The algorithm is outlined below.

*Doob-Gillespie Algorithm for drawing a single trajectory from the MPP:*

1. Set parameters $t = 0, i = 0, y_0 = (y_1, y_2, \dots, y_n)^T$ and $T = t_{max}$. Where $t_{max}$ is an upper limit on when to stop the simulation.
2. While($t < T$){

    2.1 Set $y = y_0$ .

    2.2 Record $y_{t_i}$ as being equal to $y$ and $t_i$ as being equal to $t$.

    Determine the set of possible transitions from state $y$ and record these as $y + v_1, y + v_2, \dots, y + v_m$.

    2.3 For each of the possible transitions, calculate the transition rates and record these as $q(y, y + v_1)$, $q(y, y + v_2)$, …, $q(y, y + v_m)$.

    2.4 If($\sum_{j=1}^{m} q(y, y + v_j) == 0$){

        Terminate the while loop and go to step 3.

    }

    2.5 Sample $\delta_t \sim Exponential(\sum_{j=1}^{m} q(y, y + v_j))$ and set $t$ to be equal to $t + \delta_t$.

2.6 Sample from the set of vectors $\{v_1, \ldots, v_m\}$ so that $v_k$ is selected with probability $q(y, y + v_k)/\sum_{j=1}^{m} q(y, y + v_j))$.

2.7 $i = i + 1$.

}

3. A single trajectory sampled from the MPP is then given by the collection of event times $\{t_o, \ldots, t_n\}$ and the corresponding collection of state vectors $\{y_{t_0}, \ldots, y_{t_n}\}$.

#### *A2. Conceptualising Mosquito Life Stages*

Our model is based around a simple compartmental model for the life-cycle of a mosquito which is depicted in Figure A2.1. We conceptualise a population as consisting of wildtype individuals who do not carry the *Wolbachia* bacteria and J subpopulations that each carry only a single strain of the Wolbachia bacteria (the strains are denoted *w*1, … *w*J). We assume *Wolbachia* strains *wi* and *wj* (*i* ≠*j*) are cytoplasmically incompatible. We model individuals of wildtype and infected with each of the strains using the same compartmental modelling approach as seen in Figure A2.1.

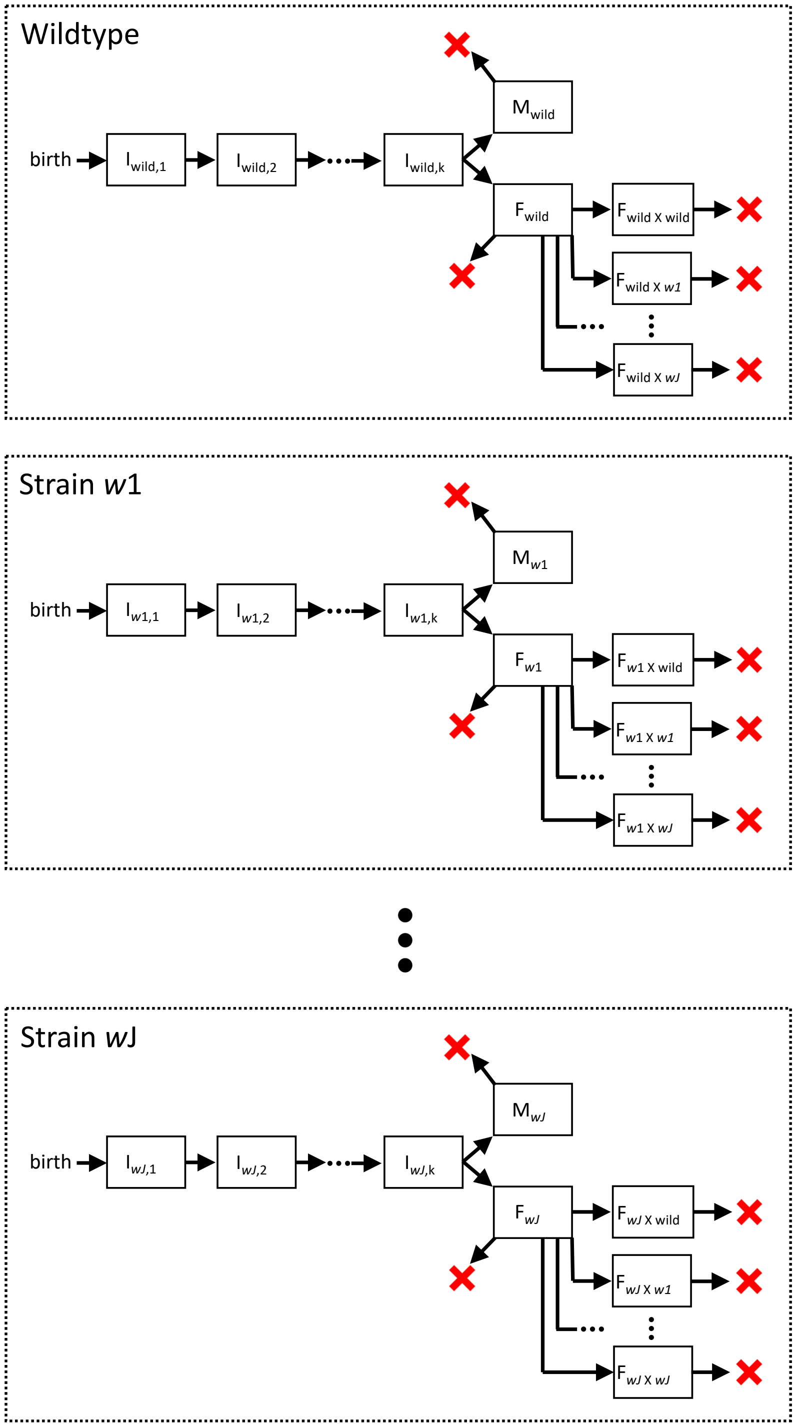


Figure A2.1. Schematic outlining the states within the MPP model. Arrows show the transitions between states that individuals can make. Arrows with red crosses show transitions to a death state for an individual.

Our model is developed to study the effectiveness of IIT population suppression strategies and is therefore largely focused on the interactions of reproducing adults. To maintain as much simplicity as possible, we avoid modelling the individual life stages of immature mosquitoes. However, immature individuals cannot be ignored, since this is the life-stage at which density dependent population regulation occurs and where an important time-lag exists between the act of egg-laying and the emergence of new adults. We adopt a parsimonious approach and choose to model a conceptual pool of immature individuals, all of which will survive to become future adults (is this similar to one of the other papers?). Simplifying the model in this fashion circumvents the need to: (i) model a large population of immature individuals, resulting in significant computational savings; and (ii) define parameters governing larval mortality. In the absence of such a simplification, we would need to incorporate a detailed reproductive model that accounted for female clutch size, egg mortality, density-dependent mortality rates and development rates for each larval stage (1st, 2nd, 3rd and 4th instars and pupae). There is another significant advantage to structuring the model in this way: only a small number of easily accessible parameters (e.g. adult carrying capacity and adult death rates) need to be specified to use it. A number of parameters that are inherent to the model (e.g. the rate at which females are mated by males and the rate at which a mated female give birth to "future adults") can be obtained from the equations that define the stable equilibrium for a purely wildtype population. In what follows, we refer to the parameters that a model user is required to specify as "primary parameters". "Secondary parameters" are those that are subsequently dictated from combining the primary parameters with our knowledge of the stable equilibrium for pure wildtype populations.

In our MPP model, the pool of immature future adults has been conceptualised as a pool of k "pseudo life-stages" connected in series and denoted $I_1, I_2, \dots, I_k$ in Figure A1. A subscript is used in these compartments to differentiate between the different subpopulations (wildtype and the *J* strains of *Wolbachia*). Our construction of these pseudo-life stages is closely related to the approach taken by Ross et al (2009) in their SI(k)R model of influenza, where an is modelled as passing through several pseudo-stages of infection. The reason for utilising this approach within the MPP is that the period of time an individual spends in a given state follows an exponential distribution. This is a consequence of the Markov property that underpins CTMCs. Exponentially distributed holding times for states is reasonable for some phenomena, such as the life-time of an adult mosquito. However, the exponential distribution has highest probability density near zero, which is not consistent with the development time of immature mosquitoes. By representing the immature phase by a series of k pseudo developmental stages, we now have a MPP where the total time spent as an immature (i.e. the total time taken to progress through the k stages) follows a phase-type distribution, which can accommodate more realistic, unimodal distributions. To maintain some simplicity with this approach, we can assume that the rate at which individuals transition through these *k* stages is homogeneous (with constant rate $\gamma$), so that the total immature development time will follow a gamma distribution with shape parameter *k* and rate parameter $\gamma$. In practice, the number of immature states (*k*) and the rate of development ($\gamma$) are chosen so as to obtain a plausible probability distribution for the delay between an egg being laid and the emergence of an adult.

It is also important to note that we do not model the sex of the future adults in the immature pools of the subpopulations. Individuals transition from the $k^{\text{th}}$ stage of the immature pool to either the adult male (*M*) or adult unmated female (*F*) state at equal rates. In other words, once an immature individual becomes an adult, we then assign it to either male or female status with equal probability. This helps to reduce the dimensionality of the model state space and aids computational efficiency.

Another feature of mosquito reproduction that we have included in our model is that mosquitoes tend to mate only once. In our model this dictates the type of offspring an individual female can produce until death. This is particularly important when we model interactions between wildtype mosquitoes and those infected with different strains of *Wolbachia*, since an incompatible mating will sterilise a female.

#### *A3. Population Dynamics and Model Transition Rates*

The MPP developed for this study consists of a state vector of length *(J + 1)(k + 3 + J)* where the element in the vector at each position corresponds to the number of individuals of that class that are currently alive. In essence, there are J + 1 subpopulations (wildtype plus each of the *Wolbachia* infected strains). A subpopulation $X \in \{wild, w1, \dots, wJ\}$, consists of the numbers of future adults in each of the *k* immature states $(I_{X,1}, I_{X,2}, \dots, I_{X,k})$, the numbers of adult males $(M_X)$, the numbers of unmated females $(F_X)$ and the numbers of females in *J+1* classes that have been mated by males from each of the subpopulations $(F_{X \times wild}, F_{X \times w1}, \dots, F_{X \times wJ})$. If we denote the numbers of individuals in each of these categories for subpopulation *X* through the vector

$$Y_X = (I_{X,1}, I_{X,2}, \dots, I_{X,k}, M_X, F_X, F_{X \times wild}, F_{X \times w1}, \dots, F_{X \times wJ})^T,$$

then the entire state vector for the MPP can be denoted as $Y = (Y_{wild}^T, Y_{w1}^T, \dots, Y_{wJ}^T)^T$.

In what follows, we define the transition rates for our MPP model. Recall that the transition rates $q(s, s^*)$ (where $s, s^* \in S$) define the probabilistic behavior of the system. For convenience moving forward, we define $z(s, "\mathrm{X}", i)$ to be a function that returns a vector as the same length as the input argument s, having zeros in all entries except at that state named "X", which takes the integer $i$ as its value. Detailed descriptions of the transition rates for the model are described under the five subheadings below.

***A3.1 Birth of an Immature Future Adult into a Subpopulation***

Density-dependent regulation of mosquito populations is believed to occur amongst larvae. As the number of immature individuals in the system increases, the chance that new offspring survive to reach adulthood, declines as a result of competition. We assume that there is a maximum number of immature future adults that can be supported in the population and denote this as $I_{max}$ the rate at which females give birth to new "future adults" is dependent upon this limiting parameter and the total number of immature future adults, $I_{total}$, in the system at a point in time:

$$q(s, s+v) = \lambda F_{wild \times wild} \frac{(I_{max} - I_{total})}{I_{max}}, \quad v = z(s, "I_{wild,1}", 1); \text{ and}$$
$$q(s, s+v) = \lambda (F_{X \times wild} + F_{X \times X}) \frac{(I_{max} - I_{total})}{I_{max}}, \quad v = z(s, "I_{X,1}", 1), \text{where}$$

$X \in \{w1, \dots, wJ\}$ and $I_{total} = \sum_{i=1}^{k} (I_{i,wild} + \sum_{j=1}^{J} I_{i,wJ})$. In essence, adult females that have undergone a compatible mating are attempting to produce new future adults at rate $\lambda$ but this rate is then moderated according to how many empty larval niches $(I_{max} - I_{total})$ are unoccupied in the population.

***A3.2 Development of Immature Future Adults in Subpopulation X***

Each individual in an immature state transitions to the next state at rate $\gamma$, so that the overall rate at which a new individual exits one of the immature states and enters the next is given by

$$q(s, s+v) = \gamma I_{X,i}, \quad v = z(s, "I_{X,i}", 1) + z(s, "I_{X,i+1}", -1).$$

***A3.3 Emergence of Adults Within a Subpopulation***

Each individual in an immature state transitions to the next state at rate gamma. For those individuals transitioning out of the final ($k^{th}$) immature state, the rate at which they enter adulthood is split evenly between the adult male state and the unmated adult female state:

$$q(s, s+v) = \frac{\gamma I_{X,k}}{2}, \quad v = z(s, "F_X", 1) + z(s, "I_{X,k}", -1); \text{ and}$$
$$q(s, s+v) = \frac{\gamma I_{X,k}}{2}, \quad v = z(s, "M_X", 1) + z(s, "I_{X,k}", -1), \text{where}$$

$X \in \{wild, w1, \dots, wJ\}$.

*A3.4 Mating of Unmated Females Within a Subpopulation*
We assume that the rate at which females are mated is independent of the density of males in the population. The transition rate is

$$q(s, s+v) = \eta F_X \frac{c_Y M_Y}{\sum_{g\in G} c_g M_g}, \quad v = z(s, "F_{X\times Y}", 1) + z(s, \mathrm{F_X}, -1),$$

where $c_g$ (> 0) is the mating competitiveness (equivalent to Fried's Index; Fried, 1971) of type $Y$ males relative to wildtype males (so that $c_{wild} = 1$); $G = \{wild, w1, \ldots, wJ\}$ ; $X \in \{wild, w1, \ldots, wJ\}$ and $\eta$ is the per capita mating rate of unmated females. Importantly, we do not assume that the rate at which females are mated is dependent upon the density of males in the population. This assumes that males are always proximal to females in the population.

*A3.5 Death of Adults Within a Subpopulation*
We assume that male individuals are dying at rate $\mu_M$ and females are dying at rate $\mu_F$. The transition rates for adult deaths within subpopulation X are written as:

$$q(s, s+v) = M_X\mu_M, \quad v = z(s, "\mathrm{M_X}", -1);$$

$$q(s, s+v) = F_X\mu_F, \quad v = z(s, "\mathrm{F_X}", -1); \text{ and}$$

$$q(s, s+v) = F_{X\times Y}\mu_F, \quad v = z(s, "F_{X\times Y}", -1), \text{where}$$

$X \in \{wild, w1, \ldots, wJ\}$.

### *A4. Derivation of Secondary Parameters from the Wildtype Equilibrium*
If we consider our population as consisting of only a single subpopulation of wildtype individuals, then there is a stable equilibrium population for our system which keeps the adult population close to its carrying capacity. Suppose that we know the values of primary parameters of our system, namely $\theta_1 = (\mu_F, \mu_M, K_{wild}, k, \gamma, p_{mated}, I_{max})^T$. As we shall see, given *t*, it is possible to determine what the values of the remaining parameters must be from the stable equilibrium of the model when it only contains a single subpopulation of wildtype individuals.

In a purely wildtype population at equilibrium, a difference in the death rates of males and females will mean that the stable numbers of each sex at equilibrium (denoted by the use of an overbar; e.g. $\bar{F}_{wild}$ and $\bar{M}_{wild}$ ) are not at parity. We start by noting that at equilibrium, the rate at which a female produces new "future females" will equal the rate at which females die, so that

$$\frac{1}{2}\lambda\bar{F}_{wild\times=wild} = \mu_F(\bar{F}_{wild} + \bar{F}_{wild\times wild}),$$

and similarly, for the production of new "future males", we have

$$\frac{1}{2}\lambda\bar{F}_{wild\times wild} = \mu_M\bar{M}_{wild}$$

Again, for clarity, the use of the bar above these variables is used to indicate the value taken at the stable equilibrium. This dictates that

$$\mu_F(\bar{F}_{wild} + \bar{F}_{wild\times wild}) = \mu_M \bar{M}_{wild},$$

so that the ratio of males to females is given by

$$\frac{\bar{F}_{wild} + \bar{F}_{wild\times wild}}{\bar{M}_{wild}} = \frac{\mu_M}{\mu_F}.$$

In other words, at a stable equilibrium population of

$$K_{wild} = \bar{M}_{wild} + \bar{F}_{wild} + \bar{F}_{wild\times wild},$$

the corresponding numbers of males and females are

$$\bar{M}_{wild} = \frac{K_{wild}}{1+\theta},$$

$$\bar{F}_{wild} = \frac{K_{wild}\theta(1 - p_{mated})}{1+\theta}, \text{ and}$$

$$\bar{F}_{wild\times wild} = \frac{p_{mated} K_{wild}\theta}{1+\theta},$$

where $\theta = \frac{\mu_M}{\mu_F}$.

For a pure wildtype population, there are three equalities that must hold for the system at the stable equilibrium:

*Equality 1.* The emergence of adults from pool $I_{wild,k}$ must equal the deaths of adults.

*Equality 2.* The births of juveniles into pool $I_{wild,1}$ must equal the emergence of adults from pool $I_{wild,k}$.

*Equality 3.* The rate at which new females emerge from $I_{wild,k}$ must balance the rates at which unmated females are mated and die.

Each of these equalities allows us to obtain one of the unknown parameters as a function of the equilibrium system state and known primary parameters, which we outline below.

#### *A4.1 Numbers of Future Adults (Juveniles) at Equilibrium*

Rearranging the first equality, allows us to obtain $\bar{I}_{wld,i}$ $(\forall\, i)$, in terms of the known parameters $K_{wild}$, $\mu_F$, $\mu_M$ and $\gamma$, and solving the first equality, then allows use to obtain $\lambda$ (the per female birth rate) in terms of entirely known quantities as well.

For Equality 1, we have

$$\gamma \bar{I}_{wild,k} = (\bar{F}_{wild} + \bar{F}_{wild\times wild})\mu_F + \bar{M}_{wild}\mu_M$$

$$= \frac{\mu_F K_{wild}\theta + \mu_M K_{wild}}{1+\theta},$$

and rearranging yields

$$\bar{I}_{wild,k} = \frac{K_{wild}(\mu_F\theta + \mu_M)}{\gamma(1+\ \theta)},$$

and since $\gamma$ is equal for each of the $k$ pools of immature future adults, it follows that

$$\bar{I}_{wild,i} = \bar{I}_{wild,k} \quad (\forall\ i).$$

#### *A4.2 Population Ceiling ($I_{max}$) for Future Adults*

Rearranging the Equality 2 allows us to obtain $I_{max}$ in terms of the other known parameters. Rearranging

$$\lambda\bar{F}_{wild\times wild}\frac{(I_{max} - \bar{I}_{total})}{I_{max}} = \gamma\bar{I}_{wild,k},$$

Yields

$$I_{max} = \frac{\bar{I}_{total}}{1 - \dfrac{\gamma\bar{I}_{wild,k}}{\lambda\bar{F}_{wild\times wild}}},$$

where

$$\bar{I}_{total} = \textstyle\sum_{i=1}^{k}\bar{I}_{i,wild}.$$

Clearly, there is a parametric constraint that in order for $I_{max}$ to be positive and larger than $\bar{I}_{total}$, we must have $\frac{\gamma\bar{I}_{wild,k}}{\lambda\bar{F}_{wild\times wild}} < 1$.

#### *A4.3 Per Capita Mating Rate of Unmated Females*

Starting with $c_{wild} = 1$, we start with Equality 3:

$$\frac{\gamma\bar{I}_{wild,k}}{2} = \mu_F\bar{F}_{wild} +\ \eta\bar{F}_{wild}\bar{M}_{wild},$$

and rearrange to obtain

$$\eta = \frac{\frac{1}{2}\gamma\bar{I}_{wild,k} - \mu_F\bar{F}_{wild}}{\bar{F}_{wild}\bar{M}_{wild}}.$$

Clearly, there is a parametric constraint that for $\eta$ to be positive, $\frac{1}{2}\gamma\bar{I}_{wild,k} > \mu_F\bar{F}_{wild}$ .

#### *A4.4 Enforcing Parametric Constraints*

When choosing primary parameters for use in the model, one must ensure that the values of the secondary parameters ensure that $\frac{\gamma\bar{I}_{wild,k}}{\lambda\bar{F}_{wild\times wild}} < 1$ and also that $\frac{1}{2}\gamma\bar{I}_{wild,k} > \mu_F\bar{F}_{wild}$, as outlined in A4.2 and A4.3. In the situation where parameters are drawn from a distribution or prior distribution, parameters that do not satisfy these constraints should be rejected as inappropriate parameter combinations.

### References


Doob, J.L. (1945). Markoff chains – denumerable case. *Transactions of the American Mathematical Society* 58(3): 455 – 473.

Fried, M. (1971). Determination of Sterile Insect Competitiveness. *Journal of Economic Entomology*, 64(4): 869 – 872.

Gillespie, D.T. (1976). A general method for numerically simulating the stochastic time evolution of coupled chemical reactions. *Journal of Computational Physics* 22(4): 403-434.

Kingman, J.F.C. (1969). Markov population processes. *Journal of Applied Probability* 6(1): 1-18.

Ross, J.V., Pagendam, D.E. and Pollett, P.K. (2009). On parameter estimation in population models II: multi-dimensional processes and transient dynamics. *Theoretical Population Biology* 75, 123-132.
