## Supplementary B for "Modelling the *Wolbachia* Incompatible Insect Technique: strategies for effective mosquito population elimination"

### *B1. Modelling Population Parameter Ranges*

Table B1.1. Primary model parameters and the lower and upper bounds of uniform prior distributions and references used to help define these.

| Parameter | Description | Units | Lower Limit | Upper Limit | References |
|---|---|---|---|---|---|
| $\mu_F$ | Per capita death rate of females | day$^{-1}$ | 0.0943 | 0.151 | Muir and Kay (1998) |
| $\mu_M$ | Per capita death rate of males | day$^{-1}$ | 0.223 | 0.562 | |
| $K_{\text{wild}}$ | Number of adults in the population at equilibrium. This parameter is sampled 20 times (to represent a suburban block with 20 houses) and the values are then summed. | (unitless) | 5 (per house) | 15 (per house) | Ritchie et al. (2013) |
| $k$ | Number of classes of future adults (integer values only) | (unitless) | 10 | 40 | Hancock et al. (2016) |
| $\gamma$ | Per capita rate of transition between future adult classes | day$^{-1}$ | 1.0 | 1.0 | Hancock et al. (2016) |
| $p_{\text{mated}}$ | Proportion of females that are in a mated state at equilibrium | (unitless) | 0.2 | 0.8 | Unpublished Field Data |
| $\lambda$ | The rate at which a single female produces future adults in an empty niche (i.e. the intrinsic rate of population growth). | day$^{-1}$ | 0.2 | 0.6 | Costero, Scott, Edman, & Clark (1998); Sowilem, Kamal & Khater (2013) |
| $c_{w\text{AlbB}}$ | The mating competitiveness coefficient (Fried's index) of *wAlbB* males relative to wild-type males. | (unitless) | 0.7 | 1.0 | Axford et al. (2016) ; Xi et al. (2005); Pagendam et al. (2018) |

### *B2. Importance Sampling Estimates of Establishment and Elimination Probabilities*

In scenarios where the FCP is very small (denoted $\beta$), we may be very unlikely to observe any simulated trajectories in which *wAlbB* establishment occurred. In such cases, the maximum-likelihood estimate of the establishment probability would be zero, despite us knowing that its value is strictly positive. We can improve our estimates of the establishment probability in such instances using a statistical method known as importance sampling (Robert & Casella, 2013). Importance sampling allows us to use simulations where the FCP was set to $\gamma$, where $\gamma > \beta$, to estimate the establishment probability for the case where the FCP is actually $\beta$. Suppose we simulate $m$ trajectories under a number of different FCPs denoted $\gamma_1, \dots, \gamma_m$. We estimate the establishment and elimination probabilities for the simulated scenario under an FCP of $\beta$ using the self-normalising importance sampling estimate:

$$\hat{p} = \frac{\sum_{i=1}^{m} \mathbb{I}_i w_i}{\sum_{i=1}^{m} w_i},$$

where: $w_i$ is the weight applied to the $i^{\text{th}}$ simulation under an FCP of $\gamma_i$; and $\mathbb{I}_i$ is an indicator variable that takes the value 1 where the corresponding simulation resulted in the event of interest (e.g. establishment or elimination) and takes the value 0 otherwise. We advocate the use of a "self-normalizing" estimator, since it ensures that our estimates of the probabilities are bounded to the interval [0, 1] and typically have smaller mean square error that the unbiased importance sampling estimator (Robert and Casella, 2013, p95). Mathematically, $w_i = \frac{\beta^{f_i}(1-\beta)^{m_i}}{\gamma_i{}^{f_i}(1-\gamma_i)^{m_i}}$, where $f_i$ and $m_i$ are the total number of females and males respectively, released in the $i^{\text{th}}$ simulation under FCP $\gamma_i$. We can also generate (1- $\alpha$)% confidence intervals as per Hesterberg (1996) using $\hat{p} \pm t_{\frac{\alpha}{2},n_e}\hat{\sigma}$, where $t_{\frac{\alpha}{2},n_e}$ is the $\frac{\alpha}{2}$ quantile of a t-distribution with $n_e$ degrees of freedom,

$$\hat{\sigma} = \left( \frac{\sum_{i=1}^{m} (\mathbb{I}_i - w_i \hat{p})^2}{m(m-1)} \right)^{1/2}$$

is the standard error and $n_e$ is the effective sample size, computed as:

$$n_e = \frac{\left(\sum_{i=1}^{m} w_i\right)^2}{\sum_{i=1}^{m} w_i{}^2}.$$

We employed this importance sampling scheme to estimate the probability of establishment and elimination under four FCPs ($10^{-4}$, $10^{-5}$, $10^{-6}$ and $10^{-7}$), under the three release strategies (constant, adaptive and crude adaptive) and across two overflooding ratios (5:1 and 15:1). Within each combination of the overflooding ratio and release strategy factors, importance sampling allowed us to reuse simulations across each specific FCP value. We used 15,000 simulations for the constant release strategies for each overflooding ratio and 1,000 simulations for the remaining release strategies at each overflooding ratio. The numbers of simulations were chosen to ensure that $n_e$ was at least 30 for each estimate produced. Each simulation used in the importance sampling was generated using an FCP that was sampled uniformly at random from the interval [$10^{-4}$, $10^{-7}$].

Table B2.1. Importance sampling estimates for *wAlbB* establishment probabilities under the 5:1 overflooding ratio. The merged cells for the number of simulations, highlights the use of the same set of simulations for estimates across a range of FCPs within each release strategy.

| Release Strategy | FCP | Number of Simulations | Effective Sample Size | Estimated Establishment Probability (and standard error) | Estimated Elimination Probability (and standard error) |
|---|---|---|---|---|---|
| Constant | 1E-4 | 15,000 | 1525 | 1.117 E-1 (5.371 E-3) | 0.8082 (2.059 E-2) |
| Constant | 1E-5 | | 2479 | 1.294 E-2 (9.710 E-4) | 0.9779 (2.003 E-2) |
| Constant | 1E-6 | | 1262 | 1.361 E-3 (1.337 E-4) | 0.9977 (2.819 E-2) |
| Constant | 1E-7 | | 1168 | 1.369 E-4 (1.386 E-5) | 0.9998 (2.930 E-2) |
| Adaptive | 1E-4 | 15,000 | 7396 | 2.754 E-2 (2.276 E-3) | 0.1395 (4.248 E-3) |
| Adaptive | 1E-5 | | 12916 | 2.976 E-3 (2.469 E-4) | 0.1422 (3.351 E-3) |
| Adaptive | 1E-6 | | 12192 | 2.988 E-4 (2.561 E-5) | 0.1427 (3.490 E-3) |
| Adaptive | 1E-7 | | 12102 | 2.988 E-5 (2.571 E-6) | 0.1427 (3.490 E-3) |
| Crude Adaptive | 1E-4 | 15,000 | 5586 | 1.659 E-2 (3.349 E-3) | 0.9711 (1.241 E-2) |
| Crude Adaptive | 1E-5 | | 11276 | 1.552 E-3 (2.188 E-4) | 0.9971 (9.484 E-3) |
| Crude Adaptive | 1E-6 | | 10099 | 1.546 E-4 (2.268 E-5) | 0.9997 (1.006 E-2) |
| Crude Adaptive | 1E-7 | | 9963 | 1.546 E-5 (2.286 E-6) | 0.99997 (1.0127 E-2) |

Table B2.2. Importance sampling estimates for *wAlbB* establishment probabilities under the 15:1 overflooding ratio. The merged cells for the number of simulations, highlights the use of the same set of simulations for estimates across a range of FCPs within each release strategy.

| Release Strategy | FCP | Number of Simulations | Effective Sample Size | Estimated Establishment Probability (and standard error) | Estimated Elimination Probability (and standard error) |
|---|---|---|---|---|---|
| Constant | 1E-4 | 15,000 | 642.7 | 0.2964 (1.794 E-2) | 0.5732 (1.924 E-2) |
| Constant | 1E-5 | | 177.9 | 3.130 E-2 (4.731E-3) | 0.9441 (6.804 E-2) |
| Constant | 1E-6 | | 45.37 | 2.884 E-3 (9.460 E-4) | 0.9946 (0.1162) |
| Constant | 1E-7 | | 35.60 | 2.884 E-4 (1.021 E-4) | 0.9995 (0.12673) |
| Adaptive | 1E-4 | 15,000 | 3604 | 9.399 E-2 (4.663 E-3) | 0.1939 (6.447 E-3) |
| Adaptive | 1E-5 | | 9281 | 9.663 E-3 (5.241 E-4) | 0.2229 (5.179 E-3) |
| Adaptive | 1E-6 | | 7699 | 9.696 E-4 (5.844 E-5) | 0.2255 (5.795 E-3) |
| Adaptive | 1E-7 | | 7527 | 9.699 E-5 (5.921 E-6) | 0.2257 (5.874 E-3) |
| Crude Adaptive | 1E-4 | 15,000 | 1278 | 5.103 E-2 (1.445 E-2) | 0.9192 (2.223 E-2) |
| Crude Adaptive | 1E-5 | | 5105 | 4.051 E-3 (3.866 E-4) | 0.9920 (1.434 E-2) |
| Crude Adaptive | 1E-6 | | 3423 | 4.140 E-4 (4.700 E-5) | 0.9992 (1.765 E-2) |
| Crude Adaptive | 1E-7 | | 3290 | 4.159 E-5 (4.812 E-6) | 0.9999 (1.808 E-2) |

#### *B3. Supplementary Figures*

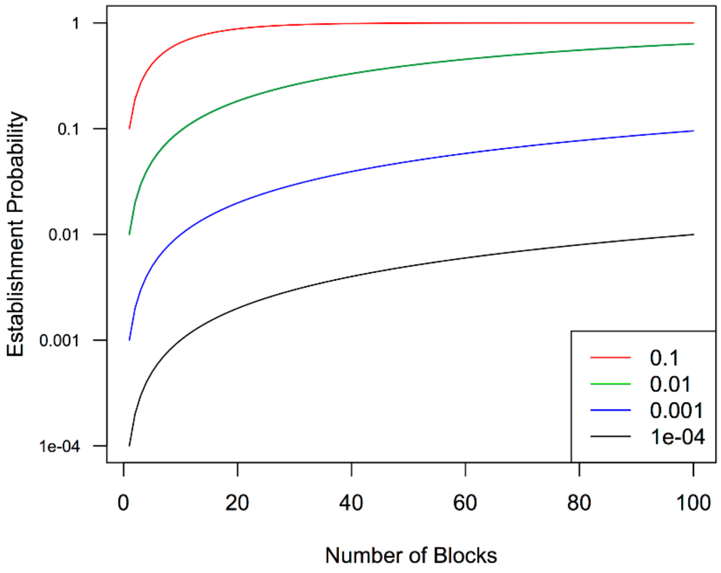


Figure B3.1 The probability of seeing one or more blocks with a *wAlbB* establishment event for increasingly large numbers of treated blocks at different block-level establishment probabilities: 0.1 (red); 0.01 (green); 0.001 (blue) and 0.0001 (black).

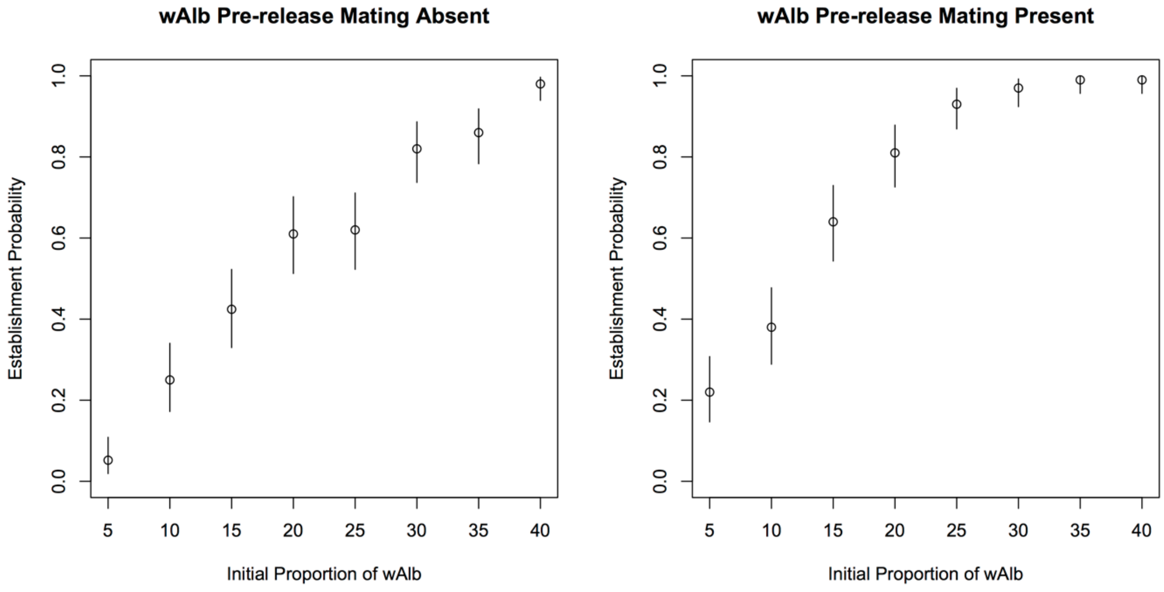


Figure B3.2. Estimates of *wAlbB* establishment probabilities across different *wAlbB* initial proportions. Each circle and vertical line shows estimated probability and 95% confidence interval (respectively) derived from 100 simulations.

### References


Hesterberg, T. C. (1996). Estimates and confidence intervals for importance sampling sensitivity analysis. *Mathematical and Computer Modelling*, 23(8), 79-85.

Robert, C., & Casella, G. (2013). *Monte Carlo statistical methods*: Springer Science & Business Media.
